# Adaptive feature detection from differential processing in parallel retinal pathways

**DOI:** 10.1101/209379

**Authors:** Yusuf Ozuysal, David B. Kastner, Stephen A. Baccus

**Affiliations:** Department of Electrical Engineering; Neuroscience Program, Stanford University, Stanford, CA; Department of Neurobiology, Stanford University, Stanford, CA, USA

## Abstract

To transmit information efficiently in a changing environment, the retina adapts to visual contrast by adjusting its gain, latency and mean response. Additionally, the temporal frequency selectivity, or bandwidth changes to encode the absolute intensity when the stimulus environment is noisy, and intensity differences when noise is low. We show that the On pathway of On-Off retinal amacrine and ganglion cells is required to change temporal bandwidth but not other adaptive properties. This remarkably specific adaptive mechanism arises from differential effects of contrast on the On and Off pathways. We analyzed a biophysical model fit only to a cell's membrane potential, and verified pharmacologically that it accurately revealed the two pathways. We conclude that changes in bandwidth arise mostly from differences in synaptic threshold in the two pathways, rather than differences in synaptic release dynamics. Different efficient codes are selected by different thresholds in two independently adapting neural pathways.

## INTRODUCTION

Sensory systems have the task of encoding a set of stimuli in a changing environment. Neural circuits meet this challenge by changing their neural code in a number of ways that can be explained theoretically by principles of efficient coding. Many sensory environments are composed of strong signals, including high luminance, high contrast, fast velocity or loud auditory stimuli. In each of these cases, in order to use a cell’s dynamic range more efficiently, sensory neurons adapt to the strong stimulus in multiple ways (1–4).

Four distinct changes in the neural code during contrast adaptation have been described in sensory neurons, including in the vertebrate retina, fly visual system and the auditory cortex (2, 5–9). Adaptation to stimulus variance changes the sensitivity, the mean response level or offset, the delay of the response, and finally the temporal frequency preference, or bandwidth. With respect to temporal bandwidth, in natural signals, low spatial and temporal frequencies are predominant, and thus nearby points in space and time have similar intensity. As such, when the the signal to noise ratio (SNR) is low, it becomes more efficient to discard weaker and noisier high frequency signals, integrating over the noise across a larger interval of space or time. At high SNR, however, cells can take advantage of the less noisy environment to reduce correlations in the input, and thus the temporal response becomes faster and more differentiating (2, 5–9). This more differentiating response preferentially encodes changes in intensity instead of the absolute intensity. These principles also account for changing spatial receptive fields, explaining why during low luminance, receptive field centers are larger and surrounds are weaker (5, 9–12). A change in temporal bandwidth during contrast adaptation has been observed in the vertebrate retina (1, 7, 13), and also in human perception (14). Although this process has been described quantitatively, and its functional importance is understood, how this change in temporal bandwidth is generated within the retinal circuit is not well described.

One obstacle to understanding the sources of these adaptive properties is that they involve nonlinear dynamic properties of the system. A second complication is that signals in the retina are merged through multiple neural pathways, each potentially with its own adaptive properties. A third challenge is that multiple changes in the neural code occur together, making it difficult to understand if any of the changes have specific mechanisms not common to other changes in encoding. Thus is it difficult to gain insights into the interactions of neural components without the use of a computational model.

Depletion of synaptic vesicles is thought to be a key source of contrast adaptation, both in terms of a change in gain, offset (15, 16) and temporal processing (17, 18). It is unknown however, how parallel pathways interact to influence any of the four adaptive properties.

Here we analyze On-Off amacrine and ganglion cells, which we find to have strong adaptive changes in their preferred temporal feature with contrast. We report a specfic dissociation between the four adaptive properties that is produced by the On pathway, in that blocking the On pathway abolishes the shift in temporal bandwidth, but leaves changes in gain, offset and the speed of response virtually intact. Our analysis first concludes that the shift in bandwidth with contrast arises from a changing ratio of activation of the On and Off pathway. Thus the temporal derivative that is computed during during high contrast is largely a difference between two neural pathways.

To understand the source of this differential activation of neural pathways in the intact system without pharmacological manipulation, we analyzed a previously reported computational model that captures all adaptive properties of On-Off ganglion cells to changing contrast (18). This model consists of a linear temporal filter, a time-independent or static nonlinearity that applies a threshold and saturation to the input, and a first-order kinetic system that creates the dynamic adaptive changes in response to a changing input (19, 20). This linear-nonlinear-kinetic model (LNK) captures all adaptive properties in an environment of changing contrast as well as the membrane potential response nearly to within the variability of the cell. Although other models have captured the properties of fast adaptation in ganglion cell membrane potential or firing rate (21–23), these models do not include the slow adaptation seen in ganglion cells, and thus would not capture all adaptive properties observed here. Given the known properties of the retina, in the LNK model, the nonlinearity corresponds to the threshold at the bipolar cell synaptic terminal, and the kinetic system captures the dynamics of synaptic vesicle release (18).

To assess why the two pathways adapted differently to contrast, we analyzed differences in the computational components of the two pathways. One might expect that because adpative properties have been assigned to the kinetics block that differential adaptation must come from different adpative kinetics in the On and Off pathwayws, representing differential properties of vesicle release. However, we find that the different thresholds in the two pathways are the primary cause of the change from the more integrating to more differentiating response, rather than the dynamics of synaptic release as has previously been proposed. These results reveal a picture where different components of adaptation are produced by different mechanisms. At the level of the ganglion cell membrane potential, changes in gain, offset and the speed of temporal processing are produced by synaptic depression (18). But to create the additional change from a more integrating to differentiating response, the differences in threshold in the two pathways leads to different levels of adaptation in the two pathways. The overall result is that increase contrast changes the mixture of the two pathways. By analyzing a model whose computational components can be mapped to neural components, different rules of efficient coding can be assigned to different internal computations and mechanisms in a parallel neural system.

## RESULTS

A randomly flickering visual stimulus was presented from a standard video monitor to the isolated salamander retina. The stimulus was spatially uniform and white in color with an intensity that changed every 30 ms, and was drawn from a Gaussian distribution with a constant mean. Every 20 s, the temporal contrast changed to one of 15 contrasts by varying the standard deviation of the distribution. To measure the total intact input to each cell, we recorded the intracellular membrane potential response from On-Off amacrine and ganglion cells using sharp microelectrodes for at least 300 s (Figure 1A). Spikes were digitally removed for analysis of the subthreshold membrane potential (18).

**Figure 1.**
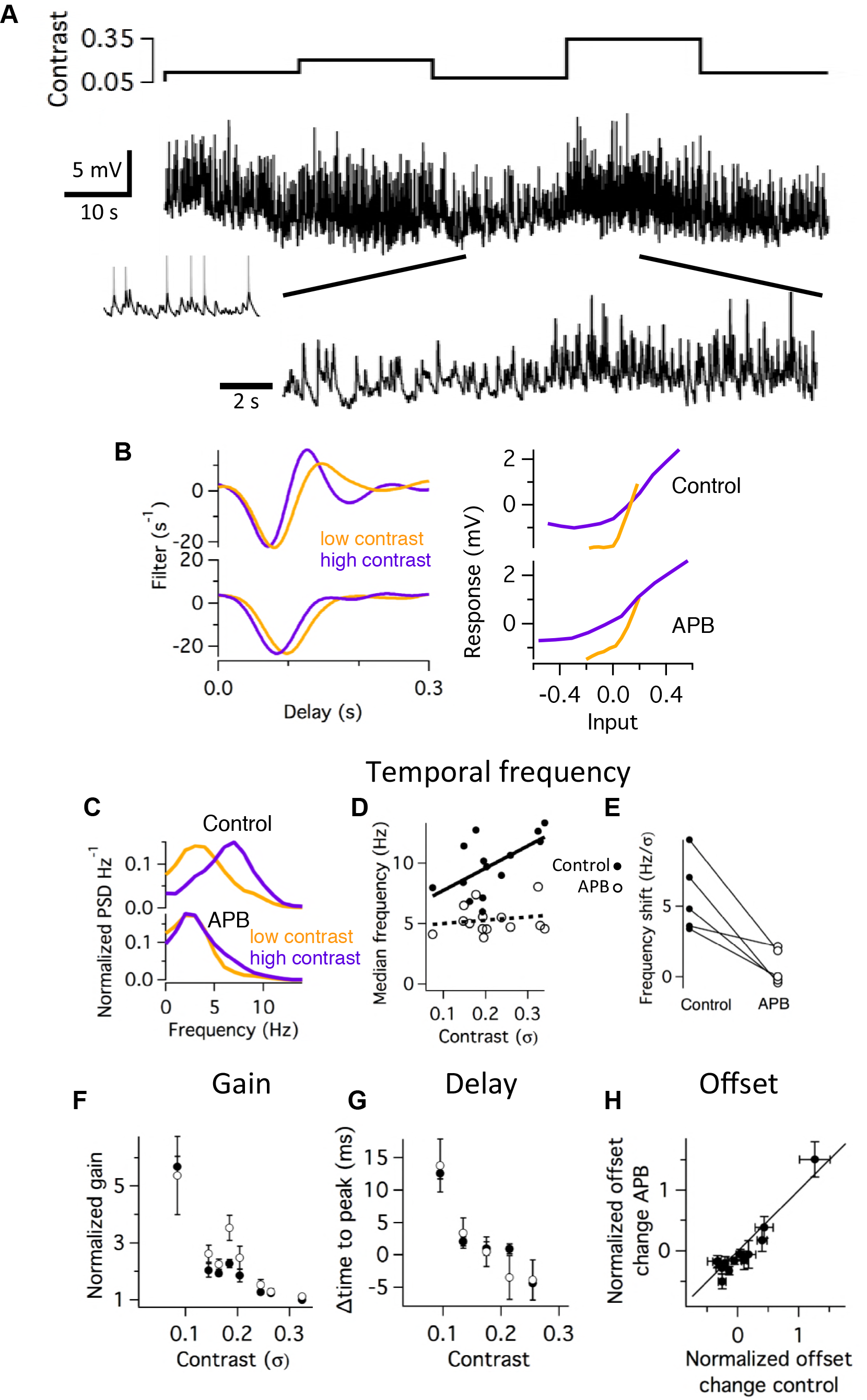
The On pathway is specifically required for temporal bandwidth adaptation. **A.** Top. Contrast of a randomly flickering stimulus drawn from a Gaussian intensity distribution with a constant mean. Contrast values ranging from 3 – 35 % were presented to the retina for periods of 20 s. Middle. Example membrane potential recording lasting 100 s of a ganglion cell responding to the 5 different contrast levels shown. Spikes were digitally removed (inset shows segment of raw recording). Bottom. Expanded segment showing transition between higher and lower contrast. **B.** Linear filters (Left) and nonlinearities (Right) fit during low and high contrast to membrane potential recordings in control solution (top) and with 30 μM APB (bottom). Filter amplitudes are normalized, with gain represented in the slope of the nonlinearity. **C.** Temporal power spectrum of the response at high and low contrasts in the control (top) and APB (bottom) conditions for an example cell. **D.** Median frequency computed as the 24) half-maximal value of the cumulative distribution function of the power spectrum as a function of contrast in the control and APB conditions for one cell. Lines indicate a linear fit. **E.** Shift in the median temporal frequency as a function of contrast, computed as the slope of the line in (**D**) for five cells, shown in the control and APB conditions. **F.** Gain of cells as a function of contrast, measured as the average slope of the nonlinearity, averaged over 5 cells. **G.** Delay in response, measured as the time to the first negative peak of the filter as a function of contrast. **H.** Offset in response measured as the mean membrane potential in 4 s segments, computed during all contrasts and compared between the control and APB conditions.

To measure the influence of the On neural pathway to contrast adaptation, we presented the stimulus in a control condition, and while blocking the On pathway using L-AP4 (APB), a metabotropic glutamate receptor agonist that blocks synaptic input to On bipolar cells. To quantify different adaptive properties at different contrasts, we used the standard approach of computing a linear nonlinear (LN) model, consisting of a temporal filter that represents the average feature conveyed by a neuron, and a static, or time-independent nonlinear function that captures threshold, gain and any saturation (7). We examined the change in temporal bandwidth produced by contrast adaptation by examining the linear filter. As previously reported (7, 24), the filter at low contrast was more monophasic, and at high contrast the filter was more biphasic (Fig.). We quantified this effect by examining the filter in the frequency domain at each contrast. The median temporal frequency of the filter was lower at low than at high contrast. In addition, the power at the lowest frequency was greatly attenuated at high contrast, reflecting the more biphasic filter (Fig. 1 F - H). We computed the average shift in the median frequency as a function of contrast, which was 5.7 ± 1.2 Hz/σ (n = 5), where σ is contrast units. In APB, the filter changed its temporal frequency much less, shifting 0.7 ± 0.5 Hz/σ, which was significantly less than the control condition (P < 0.02).

Examining other adaptive properties, we found that the average gain of a cell, represented by the slope of the nonlinearity, decreased with contrast both in the control and APB conditions, with little difference in the two conditions (Fig. 1B, C). We quantified the speed of the response as the delay until the trough of the linear filter, which changed with contrast as expected. The contrast-dependent change in the delay of the response also was not influenced by APB (Fig. 1D). We further compared the offset of the response by computing the mean membrane potential at different times during the recording, and compared them in the control and APB conditions. The On pathway was similarly not required for adaptive changes in the mean membrane potential (Fig. 1E). In total, the effect of the On pathway on adaptation was highly specific, in that only the adaptive shift in temporal bandwidth required both On and Off pathways, whereas changes in gain, response delay and offset did not.

### On-Off cells have greater changes in temporal bandwidth

We then recorded the spiking responses of ganglion cells with a multielectrode array in response to a spatiotemporal stimulus in a control condition and with APB. Cells included both On/Off cells and cells with no On pathway input. We first identified the relative strength of the On and Off pathways using a uniform field flash, and computed the ratio of the firing rate of the On and Off responses (Fig. 2A). Then we presented a one-dimensional spatiotemporal stimulus consisting of randomly flickering lines drawn from a Gaussian distribution that alternated between high contrast (4 s) and low contrast (16 s) (Fig. 2B). LN models during high and low contrast consisted of a spatiotemporal filter calculated as the spike-triggered average stimulus, followed by a static nonlinearity computed after convolving the entire spatiotemporal filter with the stimulus (25). To analyze the time course of the response, we extracted the average time course of the spatiotemporal filter as the first principal component of the temporal dimension of the spatiotemporal map (Figure 2 – figure supplement 1).

**Figure 2.**
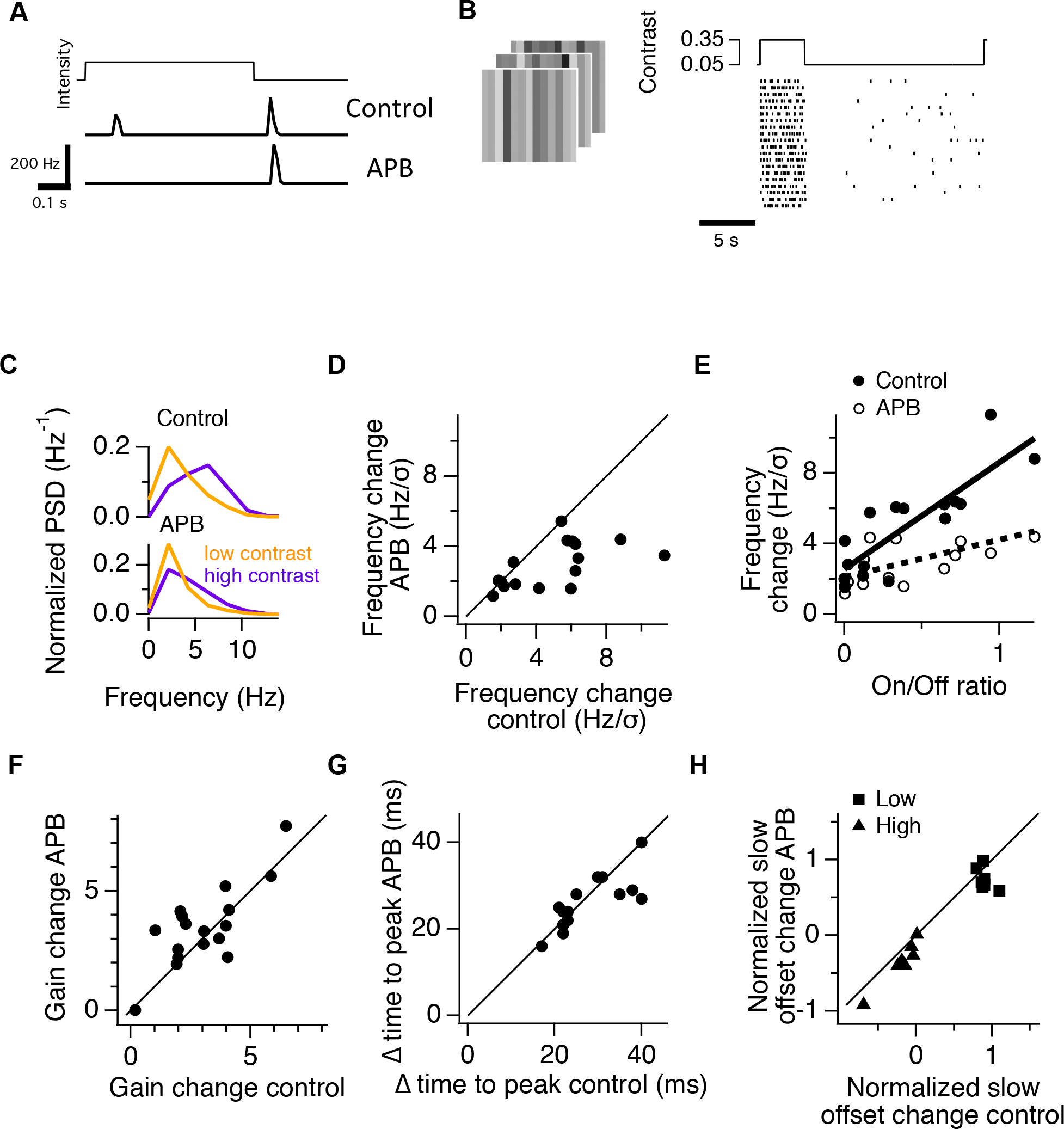
Cells with more On pathway input have a greater shift in temporal frequency. **A.** Peristimulus time histogram of the spiking response of an example On-Off ganglion cell recorded with a multielectrode array, with and without the application of APB. **B.** Example spiking response of a ganglion cell during a one-dimensional spatiotemporal white noise stimulus that alternated between high (35 %) contrast for 4 s, and low (5 %) contrast for 16 s. In the raster plot, each line is the response for an example cell to a different white noise sequence. **C.** Power spectrum of temporal filters of an example cell for high and low contrast in a control condition (top) and with APB (bottom). The LN model for the cell is shown in (Figure 2 – Figure Supplement 1). **D.** The shift in the median temporal frequency of the filter during a change in contrast compared for the control condition and in APB. The shift in temporal frequency is shown divided by the contrast shift of 30 %, (from 35 % to 5 %) to compare with the results of Fig. 1. **E.** The change in frequency shown as a function of the On/Off ratio of each ganglion cell for a population of 16 cells. Also shown is this frequency change for the same cells in APB. Lines are linear fits to the data. **F.** Gain change computed as the change in the average slope of the nonlinearity between high and low contrast compared with and without APB. **G.** The change in the time of the first negative peak of the linear filter between high and low contrast, with and without APB. **H.** The slow change in average firing rate computed from a linear fit to the firing rate during either low contrast (▪) or during high contrast (▴). Changes in rate were normalized by the maximum rate in that interval. Data was only analyzed from cells that adapted during low contrast (n = 7), increasing their firing rate. Cells that sensitized, decreasing their firing rate during low contrast (53) were excluded.

As in the case of membrane potential recordings, high contrast shifted the temporal bandwidth to higher temporal frequencies (Fig. 2C - E). Across all cells, APB reduced the shift in frequency from 5.0 ± 0.7 to 2.9 ± 0.3 Hz/σ (n = 16, P < 0.01, paired t-test) for the 0.3 σ change in contrast (Fig. 2D - E). In addition, we found that cells with the largest changes in temporal bandwidth had the greatest input from the On pathway. Cells with the highest On/Off ratio changed their temporal bandwidth by a factor of 3.6 ± 1.3 (standard bootstrap) greater than cells with the smallest On/Off ratio, computed from a linear fit to the population (Fig. 2E). As with membrane potential recordings, APB had little effect on adaptive changes in gain, delay or offset with contrast (Fig. 2F - H). These properties did not change substantially with On/Off ratio (Figure 2 – Figure supplement 2), with the exception of slow changes in offset during high contrast, which decreased for cells with greater On/Off ratio. However, APB had little effect on this relationship.

We further analyzed bipolar cell recordings with the expectation that since they receive input from a single pathway, they would not have large changes in temporal frequency with contrast. Consistent with this idea, although bipolar cells showed changes in gain that are smaller than that of ganglion cells, as previously reported (7, 26), they did not change their temporal frequency with contrast (Figure 2 – Figure Supplement 3).

From both intracellular membrane potential recordings and extracellular recordings, we conclude that cells with strong On and Off input have the highest shift in temporal frequency during contrast adaptation. Furthermore, blocking the On pathway specifically reduces this shift in temporal frequency but not the adaptive properties of changes in gain, delay or offset.

### LNK model captures the separate contribution of On and Off pathways

To consider how one can interpret our pharmacological results, in general, the response of a cell is *F*(*B*_*ON*_)+*G*(*B*_*OFF*_)+*H*(*B*_*ON*_, *B*_*OFF*_), where *F*(*B*_*ON*_) and *G*(*B*_*OFF*_) are the additive contribution from depolarizing and hyperpolarizing bipolar cells, respectively, and *H*(*B*_*ON*_, *B*_*OFF*_) is the contribution from nonlinear interactions between the two pathways. The application of APB reveals input from *G*(*B*_*OFF*_) alone, subtracting both *F*( ) and *H*( ). If there were no interactions between the two pathways (*H* ( ) = 0), then APB could reveal the separate contributions of both Off and On pathways, *F*( ) and *G*( ). In the intact circuit, however there are many potential nonlinear interactions that could occur between On and Off pathways, as numerous types of amacrine cells deliver inhibition from one pathway to another, known as ‘crossover inhibition’ (27). Thus one cannot without further knowledge simply assume that the two pathways are independent, and subtract the Off pathway under APB from the total response to discover the separate contribution of the ON pathway. However, this possible approach is indicated by the success of a linear-nonlinear-kinetic (LNK) model (18) (described below) with two independent pathways that we used previously to accurately capture the responses of On-Off ganglion and amacrine cells. Like most models of parallel pathways, however, it is difficult to know whether the separate model pathways correspond to separate neural pathways. We thus tested pharmacologically whether the model pathways did in fact capture the separate contributions of the On and Off pathways.

In each of the two pathways of the LNK model, the first stage consists of a linear temporal filter *F*_*LNK*_, which represents the average response at this intermediate stage to a brief flash of light. Although these filters were not constrained to have opposite polarity, one pathway contained a filter with a negative first peak, and the other’s filter had a more delayed positive first peak, apparently corresponding to the Off and On pathways, respectively (Fig, 3A, top).. Note that this filter, *F*_*LNK*_ is different from the linear filter, *F_LN_*, of an LN model, as *F*_*LN*_ contains all temporal processing for the entire system as opposed to the three part system of the LNK model where dynamics are contributed both by the initial filter *F*_*LNK*_ and the final kinetics stage. Thus the linear filter *F_LN_* is only an approximation to more complex nonlinear dynamics. In the next stage, a static nonlinearity *N*_*LNK*_ applies a threshold and saturation to the response, as well as setting a baseline sensitivity. The final stage consists of a 4-state first order kinetics model, a system that transitions between different states governed by a set of rate constants (28, 29). The output of the nonlinearity scales one or two of the rate constants in the kinetics block. For models fit to amacrine and ganglion cells, these rate constants are similar to previously measured rates of bipolar cell synaptic release (30–32). In each pathway, the output of the kinetics block is the active state. The final output is the sum of the two independent pathways, meaning that the effects of the two pathways combine additively.

**Figure 3.**
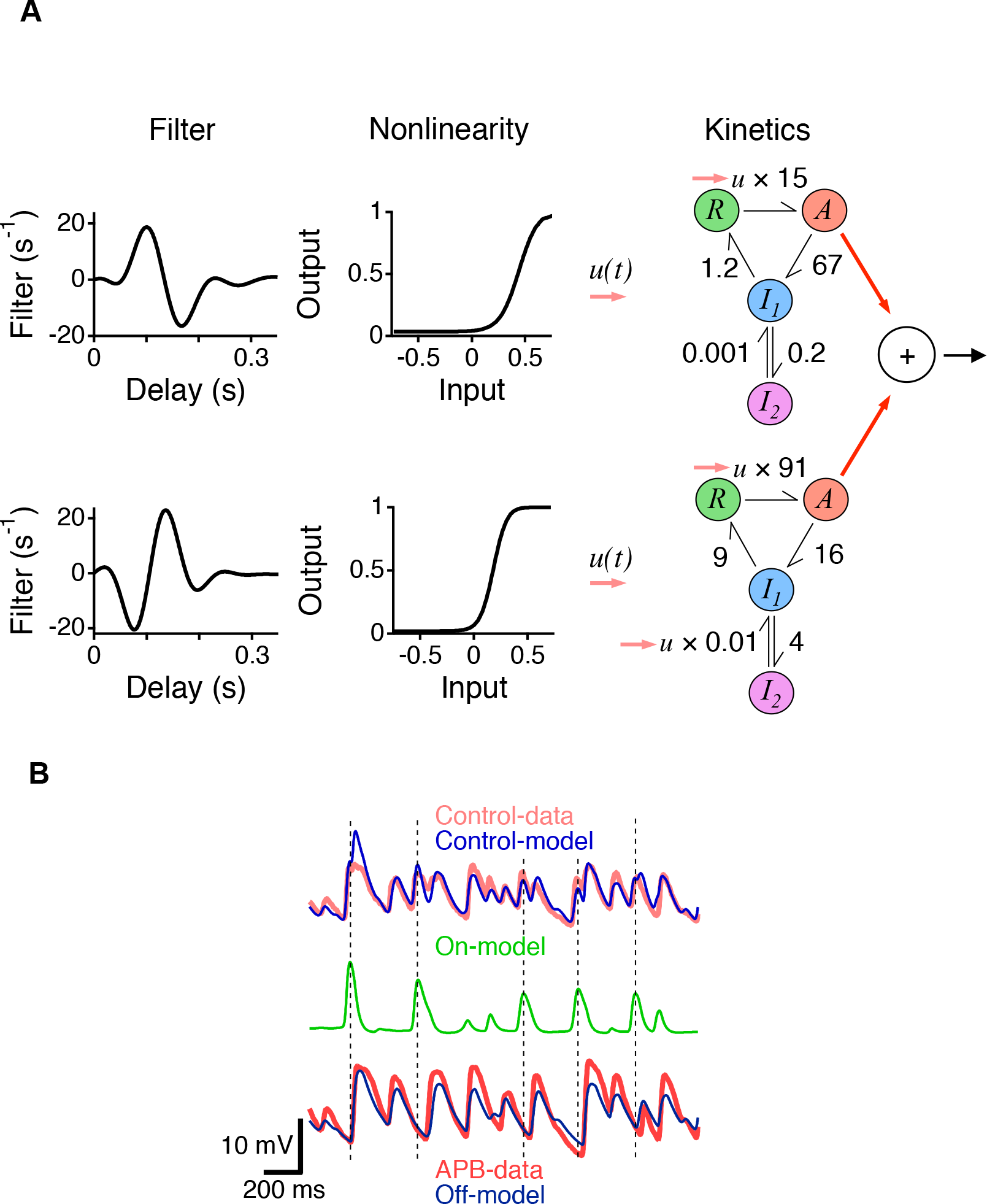
Two pathway LNK model reveals the additive contributions of the On and Off pathways. **A.** Linear – nonlinear – kinetic (LNK) model of an On-Off ganglion cell. Each of two pathways (Top, On pathway; Bottom, Off pathway), consists of a linear filter, a nonlinearity and a kinetics block. **B.** Top. Subthreshold membrane potential recordings from an On-Off amacrine cell under control conditions compared to the total output of the two-pathway LNK model. Middle. The On pathway output in the LNK model. Dotted lines indicate peaks of large responses of the On-pathway model. Bottom. Membrane potential recordings with APB compared with the Off pathway output in the control LNK model.

To verify the correspondence of the LNK model to the separate properties of the On and Off pathways, we first fit a two-pathway LNK model to membrane potential responses recorded in a control condition (Fig. 3A). The correlation coefficient between the model output and the data was (89 ± 5 % n = 4). We then blocked the On pathway pharmacologically using APB, and compared the Off pathway of the control model to a single pathway LNK model fit in the presence of APB (Fig. 3B). The correlation coefficient between the Off pathway output in the control case and the model with APB were 96 ± 1% (n = 4). Furthermore, the On pathway of the control model contained fluctuations present that were present in the data in the control case, but not when APB was present. This indicates that the drug APB, which selectively suppresses the On pathway, removed the model On pathway without changing the model Off pathway. Thus, despite the combination of two independently adapting pathways, the two-pathway LNK model accurately reveals the separate contributions from those pathways, allowing an analysis of how their components give rise to the overall behavior of the cell. Furthermore, because of the LNK model sums its two pathways, and the Off pathway fits the pharmacologically defined Off pathway, we can conclude that under these stimulus conditions the contributions of the On and Off pathways corresponded to the added parallel adaptive pathways represented by the model. Note that this does not rule out inhibitory contributions that cross pathways, but does show that the system can be analyzed as though it adds two separate adapting pathways.

### Adaptive change in stimulus feature through differential processing in two pathways

Given that under these conditions the circuit adds the two similar pharmacologically defined pathways we first analyzed the system by computing the gain of the two pathways from the recorded responses as a function of contrast. To do so, we computed LN models from the response under APB, representing the Off pathway, and from the difference between the control and APB responses, which is an estimate of the contribution that requires the On pathway. Once again, we computed the gain as the average slope of the nonlinearity. We found that at high contrast, the gain of the On and Off pathways were much more similar, whereas at low contrast, the gain of the Off pathway was much larger than that of the On pathway (Fig. 4). This analysis revealed that the two pathways adapted differently to contrast, with the Off pathway adapting more than the On pathway (Fig. 4B). Thus, as the contrast increased, different responses in the two pathways yielded a different mixture of the On and Off pathways, creating a more biphasic response than would result from a single pathway. This finding provides an explanation as to how the two pathways generate a more biphasic temporal filter at high contrast.

**Figure 4.**
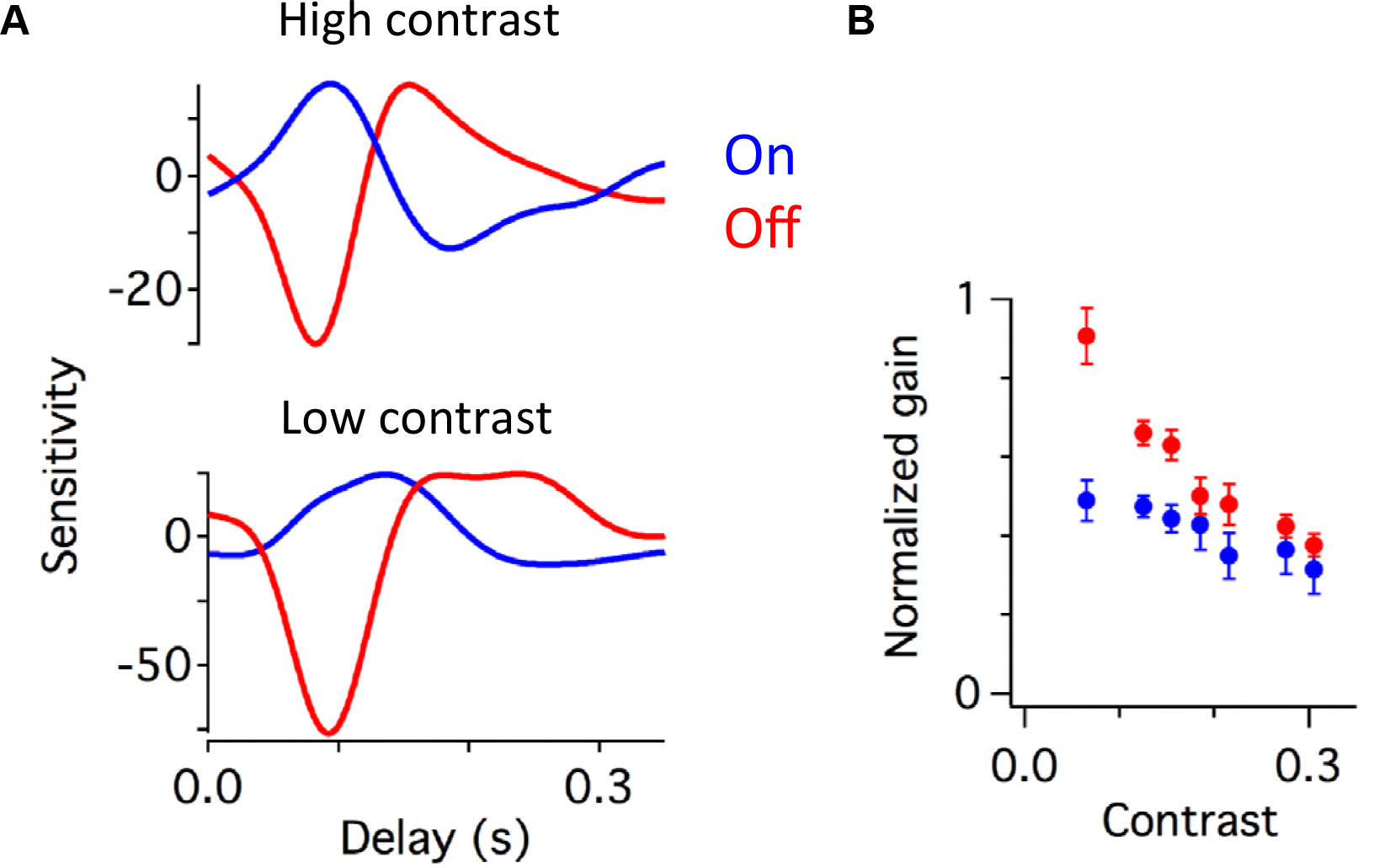
Contrast changes the weighting of On and Off pathways. **A.** Linear filters computed from the Off pathway contributions and estimated On pathway contribution at high contrast (top) and low contrast (bottom). In this representation, the gain is shown as the amplitude of the filter, so that the filter is equal to the normalized filter as shown in Fig. 1B times the average slope of the nonlinearity. The Off pathway contribution was measured as the response in the presence of APB, and the On pathway was estimated by subtracting the APB response from the control response. **B.** Normalized gain of the On and Off pathways computed as a function of contrast, measured as the amplitude of the filter in **A.** Each cell’s gain was normalized so that the total gain of the full response averaged over all contrasts was one.

### Different thresholds have a greater effect on differential processing than different adaptive kinetics

What internal computational and mechanistic properties of the two pathways underlie this differential adaptation to contrast? To answer this question, we analyzed the components of the two pathway LNK model. We first verified in the model the finding that the On and Off pathways change their relative contribution with contrast by computing the standard deviations of the output of the On and Off pathways at different contrasts. At low contrast, the On pathway contributed less to the total standard deviation of the recording (On, 22 ± 3 %, Off 78 ± 12 %, n = 8), whereas at high contrast, the contribution of the two pathways were more similar (42 ± 2 % for On and 58 ± 2 % for Off). As the contrast increased, first the output amplitude of the Off pathway increased more rapidly than that of the On pathway (Figure 5A). Then the amplitude of the Off pathway reached a plateau, which indicated that adaptive decrease in gain compensated for the increased input. Consistent with the analysis of the data using LN models (Fig. 4), the output of the On pathway increased steadily throughout the contrasts tested, adapting less than the Off pathway.

**Figure 5.**
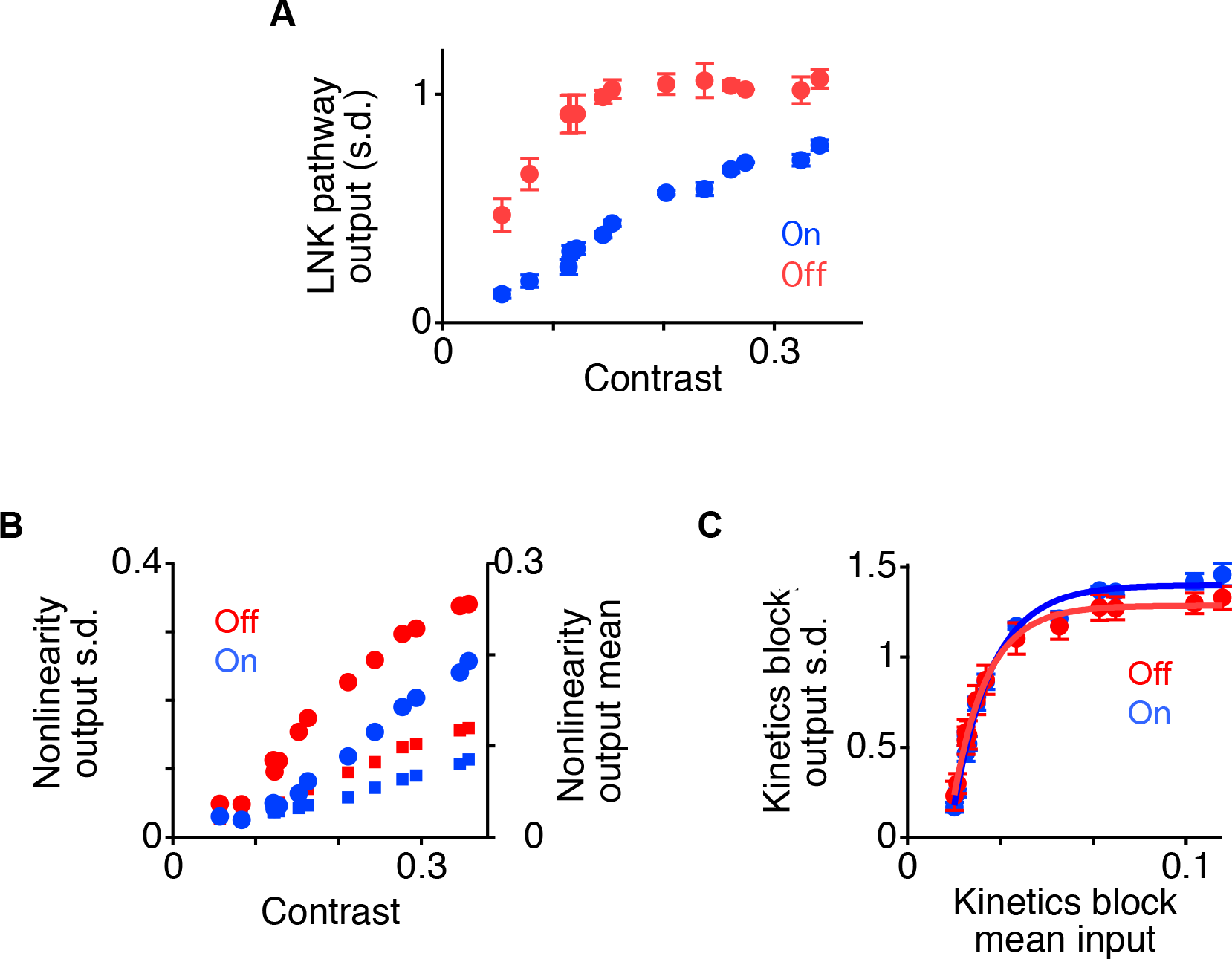
Differential adaptation in On and Off pathways. **A.** Standard deviation for the On and Off pathway outputs of LNK models of eight amacrine and ganglion cells as a function of contrast after normalizing the standard deviation of the entire output to one. Numbers do not sum to one because signals from the On and Off pathways partially cancel. **B.** Standard deviation (circles) and mean (squares) of the nonlinearity output *u*(*t*) in each pathway as a function of contrast. **C.** Standard deviation of the kinetics block output *A*(*t*) in each pathway as a function of the mean input to the kinetics block 〈*u*〉. A standard nonlinearity *N*_0_ with a threshold at zero was used to generate *u*(*t*) as a function of contrast.

One might think that because an LN model is by definition a nonadapting system, and that an LNK system shows adaptation, that differential adaptation would naturally be caused by differences in the parameters of the kinetics block in the two pathways. However, the adaptive properties of an LNK pathway are a result of both the threshold of the nonlinearity and the kinetics block (18). We thus analyzed how each stage of the model in the two pathways contributed to the differential change in output magnitude, and thus the adaptive change in temporal differentiation. Because the linear filters in the two pathways were normalized in amplitude, and the mean of the input is constant, the output amplitude of the first stage in each pathway simply reflects the contrast. Thus no differential processing occurs at this stage except for the difference in the preferred stimulus feature.

We measured how the output magnitude of the nonlinearity varied as a function of its input in the two pathways. Comparing the two nonlinearities, the On pathway had a higher threshold as previously reported for salamander On-Off ganglion cells (33), and a more shallow slope than the Off pathway as seen in On cells of the mammalian and amphibian retina (34–37) (Figure 3). Due to its lower threshold, the output of the Off nonlinearity increased at a lower contrast than in the On pathway (Figure 5B). However, as contrast increased, because of the steeper slope in the Off nonlinearity, the Off output then began to rise at a slower rate than the On pathway. In comparison, the output of the On pathway rose steadily with contrast above threshold.

To compare the independent contribution of the kinetics block in each pathway, we presented a standardized input *u*(*t*) to the two kinetics blocks, and measured in each pathway the magnitude of the output *A*(*t*), the active state. Previously it was shown that because of the threshold nonlinearity, an increase in contrast causes an increase in both the mean and standard deviation of *u*(*t*). However, only the mean 〈*u*〉 controls adaptation in the kinetics block, thereby controlling the standard deviation of the output *A*(*t*) (18). Thus we computed the amplitude of the kinetics block output as a function of the mean input 〈*u*〉. To generate *u*(*t*), we used a nonlinearity *N*_0_ with a threshold at zero for both kinetics blocks. This caused both the mean and standard deviation of *u*(*t*) to increase linearly with contrast. When 〈*u*〉 increased, the standard deviation of the kinetics block output *A*(*t*) increased quickly at first, but then rose with a decreasing rate as the kinetics block adapted (Figure 5C). Comparing the two pathways, the standard deviation of the kinetics block as a function of the mean input 〈*u*〉 was similar. The Off pathway, however, adapted slightly more than the On pathway in that, on average, the standard deviation of the On pathway rose only 1.22 ± 0.04 (n = 8) times more than the Off pathway across different mean inputs. Based on this separate analysis of the different stages of processing, differences in the output of the two pathways with contrast appeared to be caused more by the different nonlinearities than by the different kinetics blocks.

However, because of potential nonlinear interactions between different sequential stages of processing, we cannot assume that the above effects of the separate blocks will combine additively in the system. Thus we tested the source of differential adaptation in the two pathways by exchanging different components between the two pathways. We exchanged either the nonlinearities or kinetics blocks between pathways, and then measured the resulting effects on each pathway. First we switched the kinetics blocks while keeping the filters and nonlinearities fixed. We found a small change in the output of the two pathways as a function of contrast (Fig. 6A). Then we exchanged the nonlinearities and saw a much larger effect, that the change in output magnitude of the On pathway was much more similar the Off pathway in the control condition, and vice-versa.

**Figure 6.**
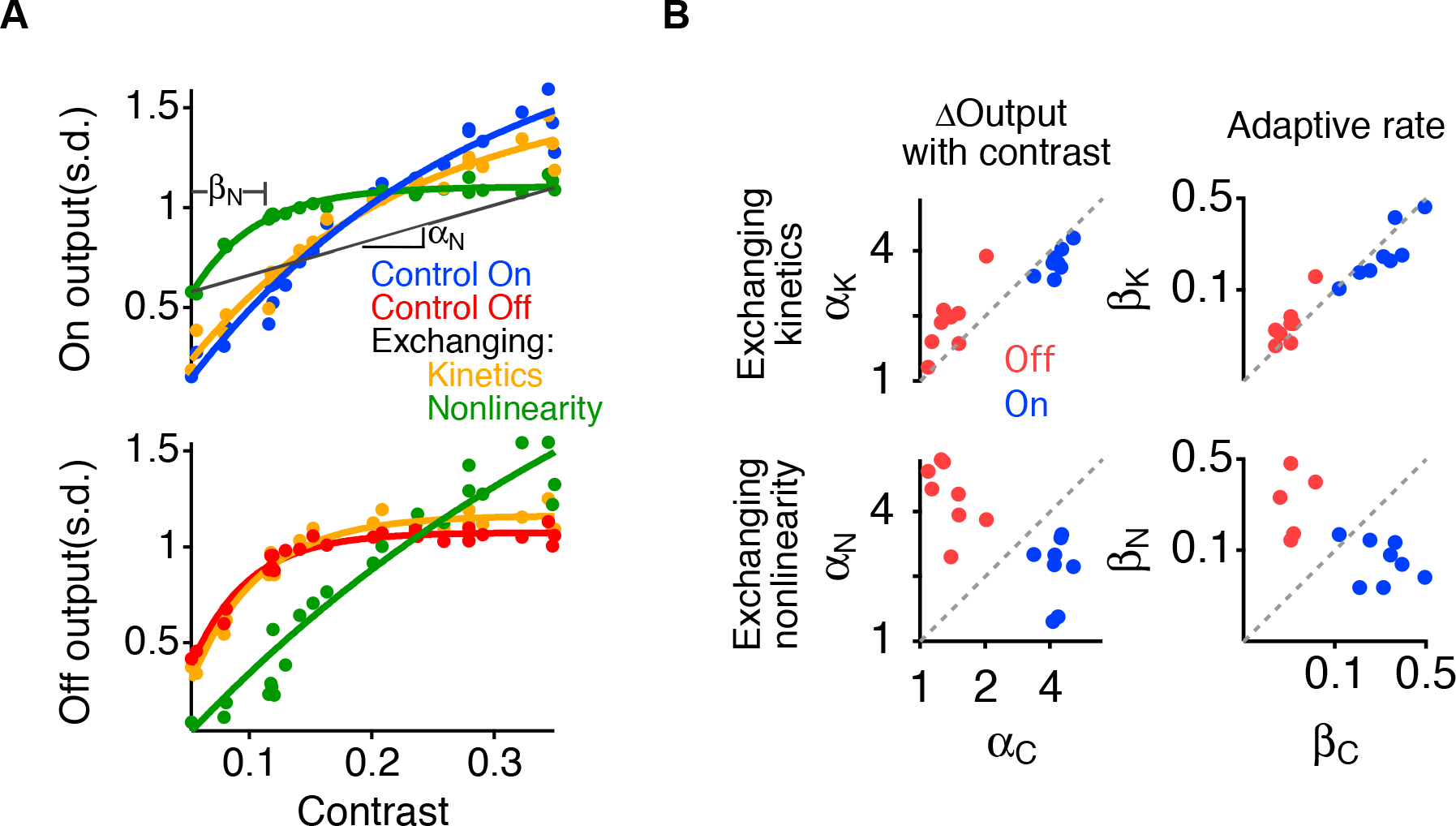
Differences in threshold create differential adaptation. **A.** Output vs contrast for the On (top) and Off (bottom) pathways of an On-Off amacrine cell, compared with the same curves after exchanging the nonlinearities or the kinetics blocks of the two pathways. Diagonal line labeled *α*_*N*_ indicates the average slope of the output vs contrast curve when the nonlinearities were exchanged. Point labeled *β*_*N*_ indicates the adaptation rate constant as a function of contrast when the nonlinearities were exchanged, computed from an exponential fit to the curve. **B.** The values for *α* (left) and β (right) after exchanging the kinetics block (*α*_*K*_, *β*_*K*_, top) or nonlinearities (*α*_*N*_, *β*_*N*_, bottom) are plotted against the values of *α*_*C*_ and *β*_*C*_ in the control condition. The scale is logarithmic.

We further quantified these effects by computing two parameters of the relationship between the contrast, *c* and output magnitude *σ* for each pathway. We computed the average slope *α* of *σ*(*c*), reflecting the average increase in output magnitude with contrast. If there was no adaptation, the standard deviation *σ*(*c*) would be proportional to the contrast with a slope of one, and so the more shallow slope of the Off pathway indicated more adaptation (Figure 6A). Then we fit an exponential function to *σ*(*c*), measuring the decay constant *β* for how fast *σ*(*c*) began to plateau with contrast. This indicates how rapidly adaptation occurred as contrast increased, with a smaller value indicating that adaptation developed at a lower contrast.

Figure 5B shows the values for the change in output *α* and rate of adaptation *β* for the two pathways in the control condition (*α*_*C*_, *β*_*C*_), when the kinetics blocks were exchanged (*α*_*K*_, *β*_*K*_) and when the nonlinearities were exchanged (*α*_*N*_, *β*_*N*_). In the control condition, the two pathways differed in the change in output *α*_*C*_ by a factor of 3.2 ± 0.2, with the On pathway having a greater slope reflecting less adaptation. Exchanging the kinetics caused the slope *α*_*K*_ to be only a factor of 1.3 ± 0.06 different than *α*_*C*_, a change of 30 %. This factor was computed by averaging *α*_*C(ON)*_/*α*_*N(ON)*_ and *α*_*K(OFF)*_/*α*_*c(OFF)*_. In comparison, changing the nonlinearities caused the slope *α*_*N*_ to be a factor 2.94 ± 0.34 different from *α*_*C*_ in the control case, a change of 194 %. Thus the change produced by exchanging the nonlinearities was 6.5 times that produced by exchanging the kinetics blocks. We then compared the control rate of adaptation *β*_*C*_ for the two pathways, which differed by a factor of 5.7 ± 0.9. Exchanging the kinetics blocks caused the rate *β*_*K*_ to be only a factor of 1.26 ± 0.07 different than *β*_*C*_, whereas exchanging the nonlinearity caused *β*_*N*_ to be a factor of 4.26 ± 0.72 different than *β*_*C*_. Thus, exchanging the nonlinearities caused a change in the adaptive rate 12.5 times greater than exchanging the kinetics. Thus, the amount of adaptive change in temporal differentiation is controlled primarily by the different nonlinearities in the two pathways, with a smaller contribution from the different adaptive kinetics. In comparison, the contrast at which the adaptive change in the temporal filter occurs is controlled almost exclusively by the differences in the two nonlinearities.

This does not suggest that synaptic kinetics plays a small role in other adaptive properties. Across contrasts, the action of the kinetics block contributes to changing the speed of the temporal filter and the gain (16, 18). These effects, however are distinct from the change in temporal differentiation controlled by differential output in the two pathways.

## DISCUSSION

Our results reveal a number of findings about contrast adaptation in On Off retinal ganglion cells, a system with multiple adaptive properties and mechanisms. First, adaptive changes in temporal bandwidth are generated in a manner distinct from changes in gain, the speed of the response and the mean response level. Second, changes intemporal bandwidth specifically require the On pathway, whereas other adaptive changes do not. Third, the mechanism that changes temporal bandwidth causes the On and Off pathways to adapt to contrast differently, changing the ratio of On to Off input at high contrast. Because the On pathway has a more delayed average temporal filter than the Off pathway, a greater On pathway contribution results in a more biphasic filter for the entire system. Fourth, we verified using pharmacology and a model that On-Off ganglion cells have the computational structure of two independent adapting pathways whose outputs are summed together. Finally, analysis of that model indicates that the key differences between the On and Off pathway that cause differential adaptation and thus underlie changes in temporal bandwidth is a difference in the threshold of the two pathways, rather than differences in their nonlinear dynamics. This implies that the differences in the threshold of the bipolar cell terminal as compared to the baseline membrane potential are responsible for the change in temporal bandwidth, as opposed to differences in the dynamics of synaptic release in the two pathways.

We have used pharmacology to block the signals traveling through one receptor type, and then infer the independent contributions of the blocked and unblocked pathways. It is important to note that the subtractive procedure we have performed to make this inference is not generally valid unless the two pathways do sum together independently. Thus, the use of the LNK model to verify this computational structure, and pharmacology to validate this aspect of the model is a necessary step in the interpretation of the two summed pathways. Crossover inhibition between the On and Off pathways is observed in many ganglion cells (27). However, our model of the Off pathway fit to the whole system captures the cell’s behavior when only the Off pathway is active, and the model’s symmetric structure lends support that blocking the On pathway can reveal the contributions of both Off and On pathways. Note that the conclusion that the cells conform to a model of two independently adapting pathways does not mean there is no crossover inhibition. The two pathways consist of one pathway that originates from APB sensitive neurotransmission, and one that does not. The APB sensitive pathway may itself be an average of several neural pathways, some that deliver a signal to some component of the Off pathway.

One reason, however, to think the effect of crossover inhibition is not strong in On-Off ganglion cells that crossover inhibition often creates a more linear cell (38), whereas On Off cells are necessarily nonlinear, and have a sharp threshold. A second effect is for the On pathway to create a more strongly rectifying Off pathway by shifting the effective threshold of the Off pathway through tonic inhibition (39). Because the same Off pathway in the model fit the response with and without the On pathway present, we predict similarly that this effect of crossover inhibition is not strong here. If there is crossover inhibition in these cells, it may contribute to the initial linear filter in each pathway, which mechanistically would correspond to presynaptic inhibition onto the bipolar cell terminal. The majority of salamander On-Off ganglion cells have direct input from both On and Off bipolar cells, with a smaller percentage (< 25 %) having input from one pathway that is delivered only via intervening amacrine cells (40). This more common type might be expected to behave according to the model of independently adapting pathways we describe here. We cannot assume, however, that On and Off pathways are strictly parallel streams that do not interact, and we expect this will change with more complex visual stimuli than the uniform field stimuli used to create the model. Nonetheless, the basic conclusion that the On pathway is required for large changes in temporal frequency but not other adaptive properties is observed with a spatiotemporal stimulus (Fig. 2).

An alternative computational structure has been proposed to account for contrast adaptation in ganglion cells using a divisive effect of one visual feature on another (22). This Div-S model can capture fast adaptation in On ganglion cells of the mouse, but does not include slow adaptive time constants and thus was not tested against the data here. The basic structure of the LNK model is a depressing synapse that follows a threshold, and thus a stronger signal changes the mean level of the input to the synapse. Because of the necessity of the threshold, this arrangement may be less effective at generating adaptation for a more linear cell. Because the Div-S model has been shown to fit On ganglion cells, which are more linear than Off cells (34–37), it may be that a divisive structure is required for adaptation in more linear cells.

### Differential adaptation in parallel pathways

Systems that can be accurately captured with a linear-nonlinear (LN) model do not change their properties of temporal filtering or sensitivity with a change in stimulus statistics. Thus, a change in the parameters of the LN model has served as a definition of adaptation (7, 24, 41, 42). Both On and Off pathways adapt independently, in that the gain of each pathway changes with contrast (18). Overall, the Off pathway adapted at a lower contrast, and reached a more complete level of adaptation at lower contrasts than the On pathway (Fig. 5A). This change in the relative strength of the On and Off pathways is consistent with an earlier study (43) that showed that one type of On-Off 19 ganglion cell – type II from that study – changed its temporal filter with contrast, but this behavior could not be reproduced by a single LNK pathway. This earlier study identified different clusters in a spike-triggered covariance analysis that could have been generated by the On and Off pathways, although the small amount of spikes occurring from the On pathway was insufficient to model. Our membrane potential recordings here provided sufficient data to fit a two pathway LNK in order to analyze the internal computational properties of the separate On and Off input. Although the different adaptive kinetics in the two pathways do make a small contribution to differential adaptation, the primary source for this adaptive change is the different thresholds in the two pathways (Fig. 5 – 6). Thus in general, changes in temporal bandwidth alone could potentially be produced through the summation of nonadapting pathways.

### Differences in On and Off inputs in On-Off ganglion cells

A number of studies in different species have shown that On and Off pathways have differences in their nonlinear properties. On ganglion cells are more linear and have lower thresholds in salamander, mouse and primate (34–37), and On pathway neurons are similarly more linear in the fly early visual system (44). Although we observe the shallower slope for the On pathway, we observe that for On-Off ganglion cells, On pathway input has a higher threshold (Fig. 3). There is substantial diversity in the responses of individual bipolar cells in salamander (40, 45) and mouse (46), and it may be that On bipolar cell inputs to On-Off ganglion cells have a higher synaptic threshold than On bipolar cell inputs to On type ganglion cells.

### Role of adaptive synapses

Properties of synaptic vesicle release have been shown to produce all types of adaptive properties discussed here. Depletion of synaptic vesicles can cause changes in gain, the elevated rate constants of release as a function of high calcium concentration can speed the overall system dynamics, and slow replenishment can create a homeostatic effect on the average response. Synaptic depression could also cause a change in temporal bandwidth, in that at high release rate, synaptic depression can create a more transient response, thus attenuate low temporal frequencies. Thus although all observed adaptive changes could potentially be carried out by synapses in a single pathway, the large shift in the temporal bandwidth is instead implemented here by a summation of two neural pathways that adapt differently to contrast. Because the effect of the dynamics of synaptic vesicle release, and the flexibility of the effect of the rate constants on adaptive behavior, one might think that the different adaptive dynamics in the On and Off pathways could arise from differences in synaptic release. Our computational analysis, however, indicates the different threshold of the On pathway, which has previously been reported, gives rise to that differential adaptation.

The LNK model breaks down the cell’s response into smaller subsystems, first by separating On and Off pathways, then further compartmentalizing computational function into feature selection, nonlinear distortion and thresholding, and adaptation. The accuracy of this model, its ability to capture adaptive properties and the correspondence of its components to distinct neural pathways (Fig. 3) allow us to localize a potentially complex response into simpler computational elements. In the model, one can think of each pathway as having a threshold for adaptation. As the variance of the signal increases, the mean of the signal after the threshold will increase. This increase in mean then triggers adaptation of the kinetic system. Thus, one can think of the threshold as being a threshold for adaptation, and it is this threshold that is different between the two pathways, although the kinetic system that implements that adaptation is similar between the pathways. The similarity of the adaptive properties is consistent with the notion that blocking On pathway does not change gain, speed or offset. If the two adaptive pathways were different one would expect that changing the mixture would change adaptation of the overall system.

Different types of mechanisms can generate changes in temporal differentiation. Photoreceptors have a more biphasic response at higher luminance (47), yet use only a single biochemical pathway to convey the light response (48). Insect photoreceptors accomplish this switch with a combination of two ionic conductances with different thresholds (49). Previous experiments give evidence as to the correspondence of different stages of the LNK model with different levels in the circuit. Because the bipolar cell membrane potential does not show strong rectification for a constant mean intensity stimulus but ganglion cells are rectified, the threshold likely arises at the bipolar cell synaptic terminal from voltage-activated calcium channels. One possible source of the different thresholds in the two pathways is differential expression of such calcium channels, which differ in rods in cones, but it is unknown whether they differ in On and Off bipolar cells (50). An alternative source is differential inhibition onto bipolar cell terminals (51), which would set the resting potential at different levels relative to the activation threshold of calcium channels in the synaptic terminal.

### Two pathways implement a change in the rules of efficient coding

The rules of efficient coding are known to change in a sophisticated way when the signal strength changes. Due to the dominance of low temporal frequencies in natural visual scenes, it can be an efficient strategy to remove these slow correlations, thus giving more equal weight to high and low frequency signals (5, 9). This approach however can come at a cost when signals are weak and noisy – rather than computing the difference between noisy signals, the more efficient strategy is not to decorrelate, but to simply exclude high frequency signals. Here we see that this shift in temporal frequency in On Off ganglion cells comes mostly from a change between one and two neural pathways.

The phenomenology that we observe with respect to a switch from one to two neural pathways is highly similar to that found in the auditory cortex, where the stimulus-response relationship changes from monophasic to biphasic at higher signal strength (**52**). In that case, there is an accompanying switch from a one-dimensional to a two-dimensional stimulus representation. However, because of a lack of knowledge of neural inputs in the cortex it is not known whether this change involves the additional contribution from a separate neural pathway as we can conclude in the retina.

Our results show that distinct properties of contrast adaptation are controlled by distinct mechanisms acting in concert, with the baseline set of properties of changing gain speed and offset present in individual neural pathways, and temporal bandwidth changing arising from the recruitment of a high threshold pathway. Thus, the system is layered, with a baseline rule of efficient coding implemented in single pathways, and a second-level rule selected or rejected by the high threshold of a second pathway. Different rules of efficient coding are selected by different combinations of neural pathways.

## Acknowledgements

We thank K. Boahen and K. Shenoy for helpful discussions, and P. Jadzinsky for comments on the manuscript. This work was supported by grants from the NEI, Pew Charitable Trusts, McKnight Endowment Fund for Neuroscience, the Alfred P. Sloan Foundation and the E. Matilda Ziegler Foundation (S.A.B.).

## Author Contributions

Y.O. and S.A.B. designed the study, Y.O. and D.B.K. performed the experiments and analysis, and Y.O. and S.A.B. wrote the manuscript.

## METHODS

Electrophysiology. For intracellular recording, the intact salamander retina of either sex was held in place under a transparent dialysis membrane containing several 150 – 300 |im holes. Intracellular electrodes were filled with 2 M potassium acetate (200 – 300 MΩ) and guided into the retina under infrared illumination viewed through a CCD camera. Bipolar cells, On-Off amacrine or ganglion cells were identified by their flash response and level in the retina. Ganglion cells were recorded in the ganglion cell layer, and On-Off amacrine cells were recorded in the inner nuclear layer. Intracellular recordings had resting membrane potentials more hyperpolarized than – 65 mV, and were made from eight On-Off cells including four ganglion cells and four amacrine cells. Results were similar for the two cell types, and are pooled in all analyses. Recordings overall ranged from 10 – 90 minutes in duration. Extracellular spiking responses from ganglion cells were recorded using an array of 60 electrodes (Multichannel Systems) as described (54).

Visual Stimulation. A spatially uniform visual stimulus was projected from a video monitor onto the retina. The stimulus intensity changed every 30 ms, and was drawn from a Gaussian probability distribution with mean intensity, *M* (~ 8 mW/m^2^) and standard deviation *W* (55). Contrast was defined as *W/M*, and changed randomly every 20 s to a value between 0.05 and 0.35, drawn from a uniform distribution. The stimulus lasted 300 s (15 contrast levels) and the identical stimulus sequence was repeated at least two times.

Standard linear model. A linear temporal filter was computed that most closely approximates the entire response of a cell or pathway. This filter *F*_*L*_(*t*) is different than the component linear filter of the LNK model, *F*_*LNK*_(*t*). Linear temporal filters were computed as described (7). The stimulus intensity *s*(*t*) was normalized to have zero mean, and a standard deviation equal to the contrast. The filter, *F_L_ (t*), was computed as the correlation between *s*(*t*) and the response *r*(*t*) normalized by the autocorrelation of the stimulus. The filter was computed as,

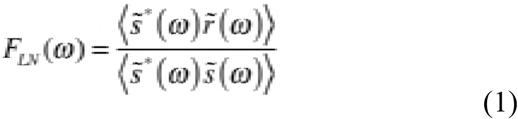

where 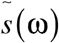 is the Fourier transform of *s*(*t*), 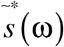 its complex conjugate, and 〈…〉 denotes averaging over 1 s segments spaced every 0.1 s throughout the recording. The denominator corrects for deviations of the video monitor from a white noise distribution (56). This calculation was performed separately at low (5 %) or high (35 %) contrast. The filter was normalized in amplitude so that when the filter was convolved with the stimulus to yield a linear prediction *g*(*t*),

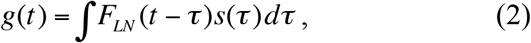

the variance of *g*(*t*) and *s*(*t*) were equal,

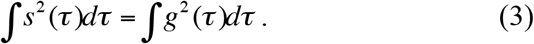

Linear Nonlinear Kinetic model. LNK models were optimized as described, (18), where additional details can be found. For On-Off cells the model had two pathways. Each pathway consisted of a linear temporal filter *F*_*LNK*_(*t*), a static nonlinearity, *N*(*g*) and a first order kinetic system defined by a transition matrix **Q**(*u*). The components were parameterized as described below, and all parameters were fit together using a constrained optimization algorithm. For each pathway, the stimulus, *s(t)*, was passed through a linear temporal filter, *F*_*LNK*_(*t*) and a static nonlinearity, *N*(*g*),

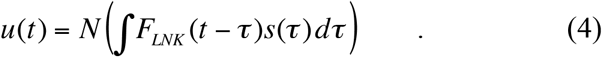

Although these two initial stages have the same structure linear-nonlinear (LN) model, the filter and nonlinearity are different functions than those computed for an LN model, and are optimized, rather than computed using reverse correlation. The kinetics block of the model is a Markov process defined by

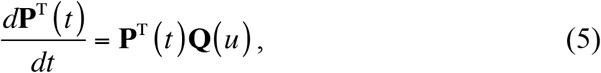

where **P**(*t*) is a column vector of *m* fractional state occupancies such that 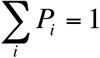, and **Q** is an *m* × *m* transition matrix containing the rate constants Q_ij_ that control the transitions between states *i* and *j*, with 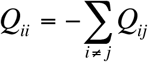. After this differential equation was solved numerically, the output of the model, *r′*(*t*) was equal to one of the state occupancies scaled to a response in millivolts,

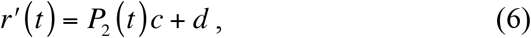

where *c* and *d* are a scaling and offset term for the entire recording.

States and rate constants are diagrammed in Fig. 2A and defined as,

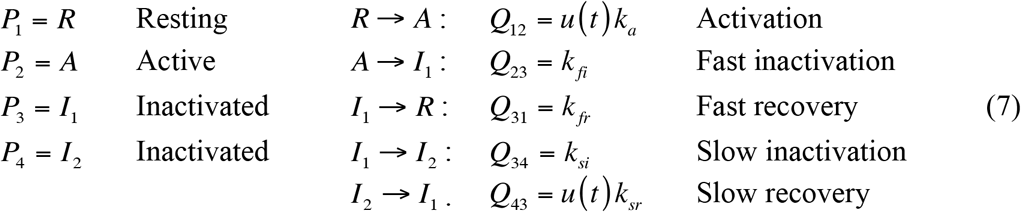

The change in state occupancy was thus determined as

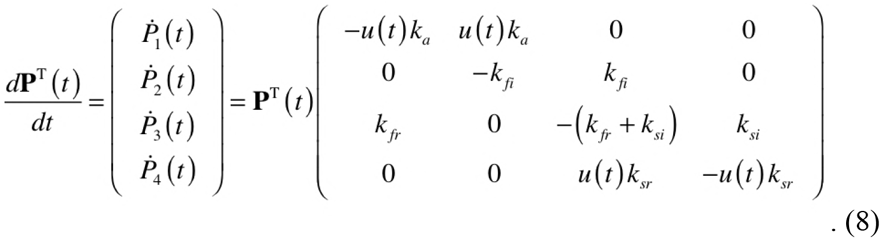

The output of the two LNK pathways were then scaled with independent weights *w*_*ON*_ and *w*_*OFF*_ and summed to give the final output of the model.

---

**Figure 1 – Figure Supplement 1.**
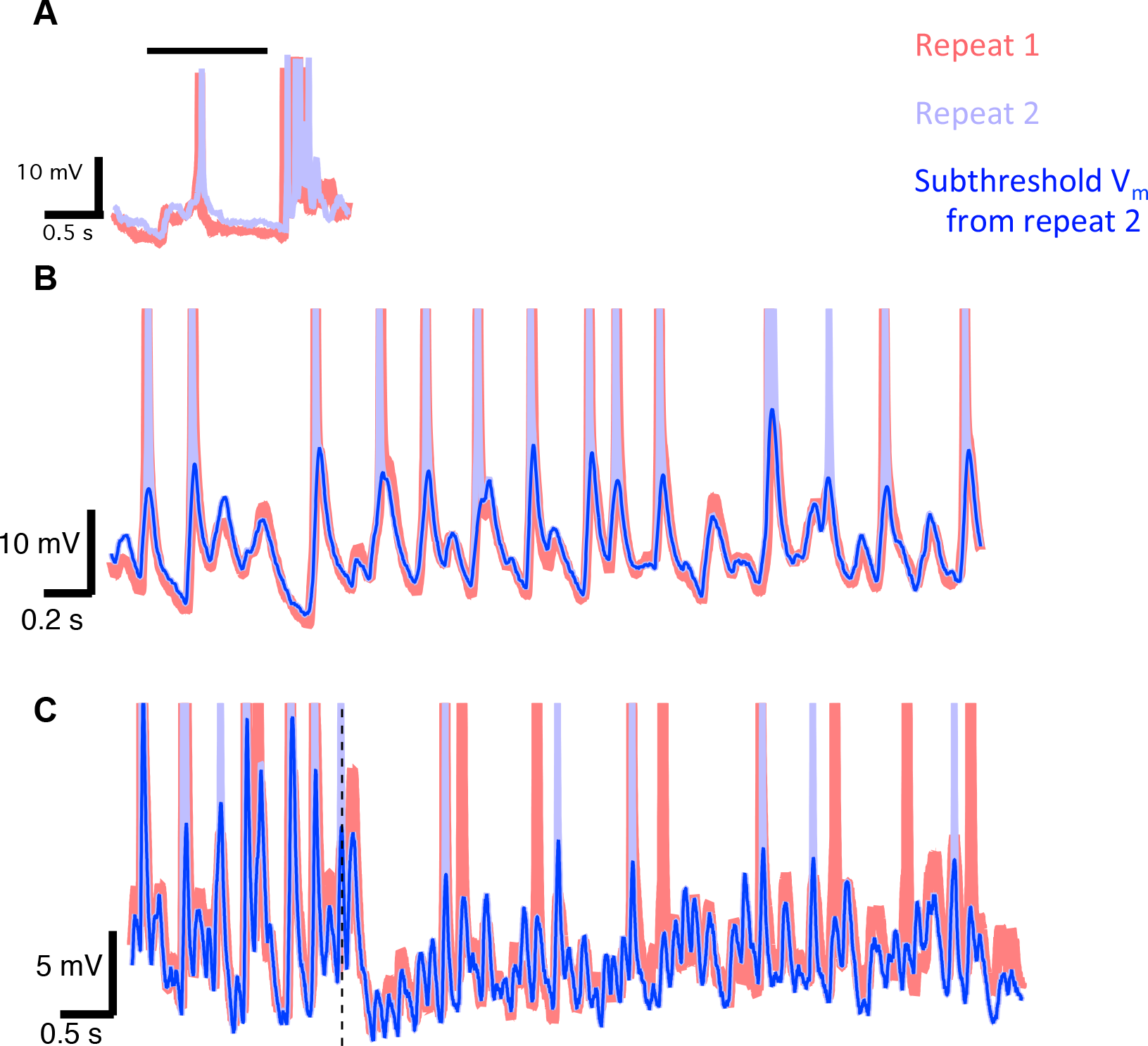
Intracellular recordings from ganglion cells at different contrasts. **A.** Two superimposed traces of the flash response from an example On-Off ganglion cell. Bar indicates the time of light On. **B.** Intracellular recording of a ganglion cell responding to two repeated presentations of the same white noise stimulus sequence consisting of a uniform field at high contrast (35 %). Also shown is the subthreshold membrane potential of one of these repeats extracted for further analysis. **C.** Same as **B** for the transition (dotted line) from high (28 %) to low (5 %) contrast. A slow recovery from hyperpolarization can be seen in the 5 % contrast traces.

**Figure 2 – Figure Supplement 1.**
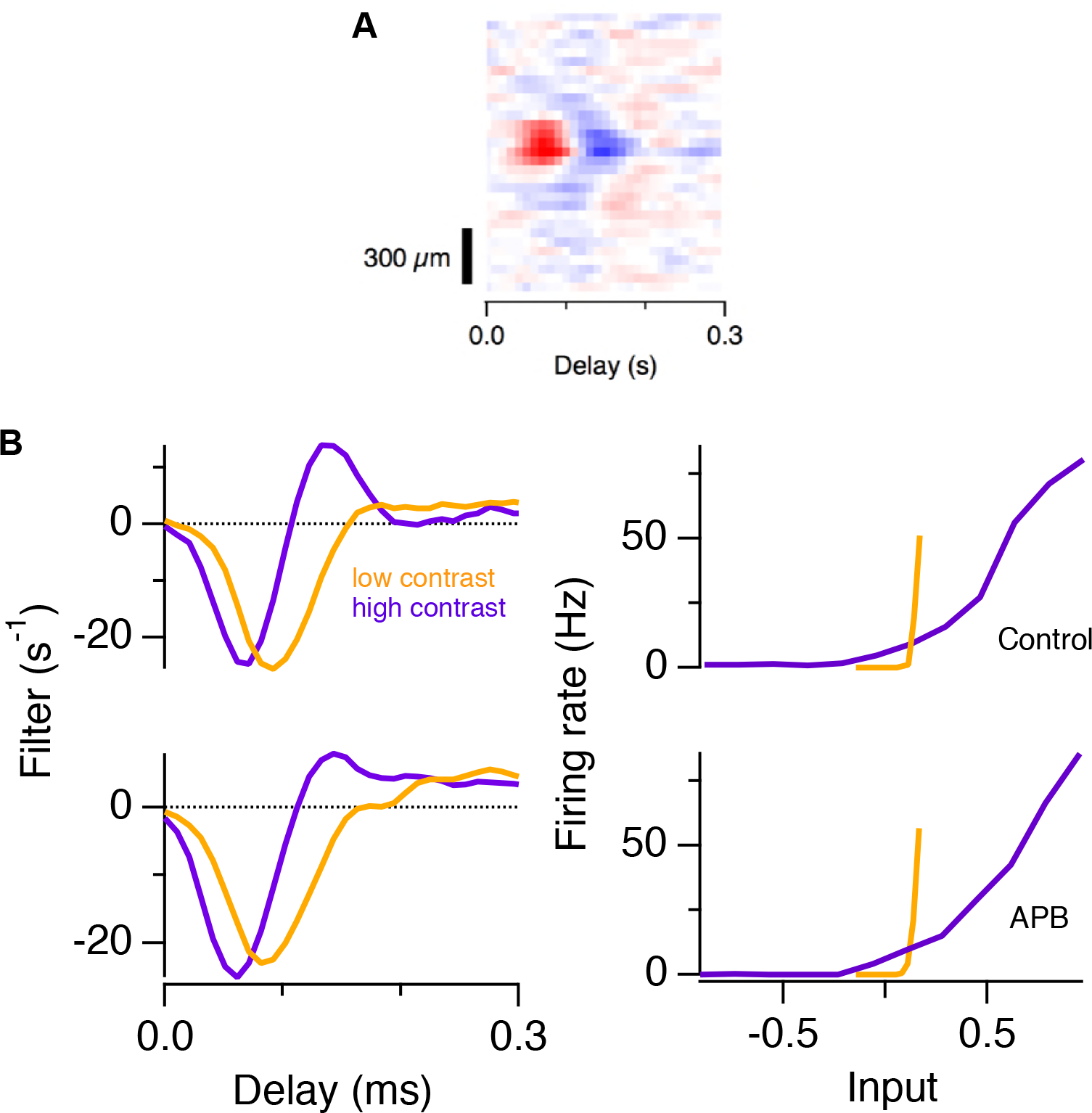
Linear-nonlinear models showing adaptation in the ganglion cell spiking response. **A.** Spatiotemporal filter of an example On-Off ganglion cell. **B.** LN models from the cell responding to high (35 %) and low (5 %) contrast in a control condition (top) and in the presence of APB (bottom). The average time course of the cell is shown computed as the first principal component of the spatiotemporal filter.

**Figure 2 – Figure Supplement 2.**
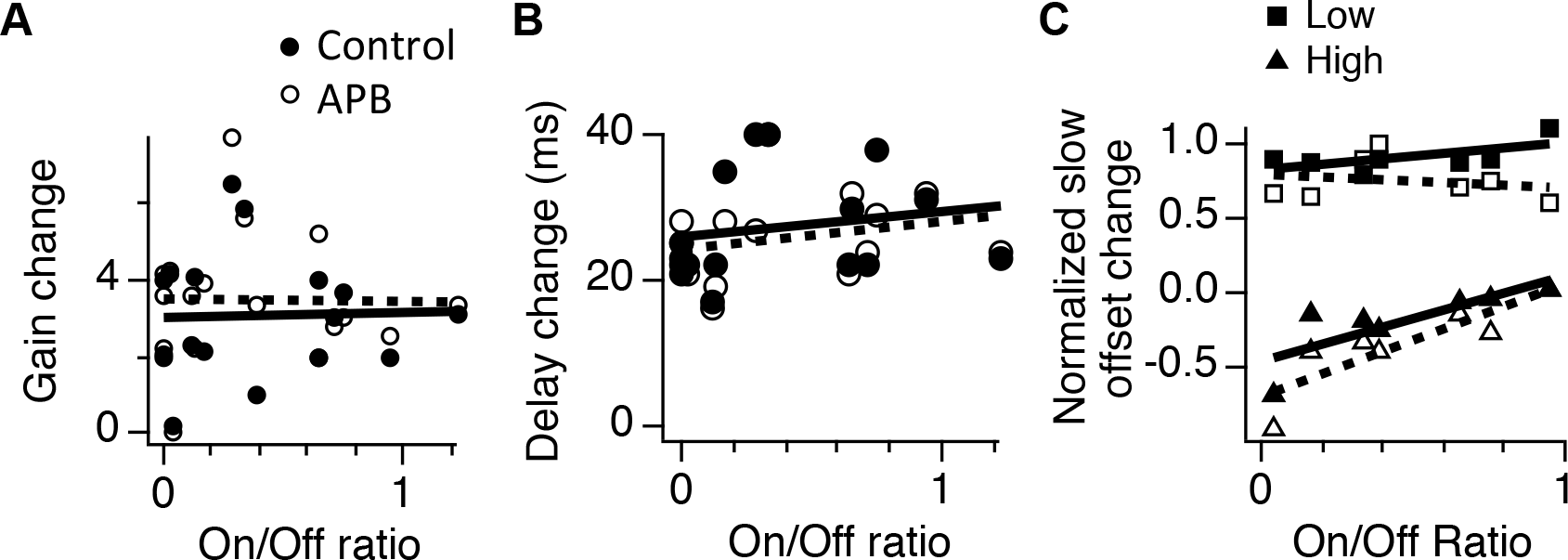
Adaptive changes in gain, delay and offset do not correlate with On pathway input. **A.** Gain change between high and low contrast for ganglion cell spiking shown as a function of the On/Off ratio of each cell. Results shown in a control condition (solid symbols and lines) and with APB (open symbols and dotted lines). Lines are fits to the data. **B.** Change in the time to the first negative peak between high and low contrast shown as a function of the On/Off ratio of each cell. **C.** Slow offset during adaptation to high or low contrast as a function of the On/Off ratio of each cell.

**Figure 2 – Figure Supplement 3.**
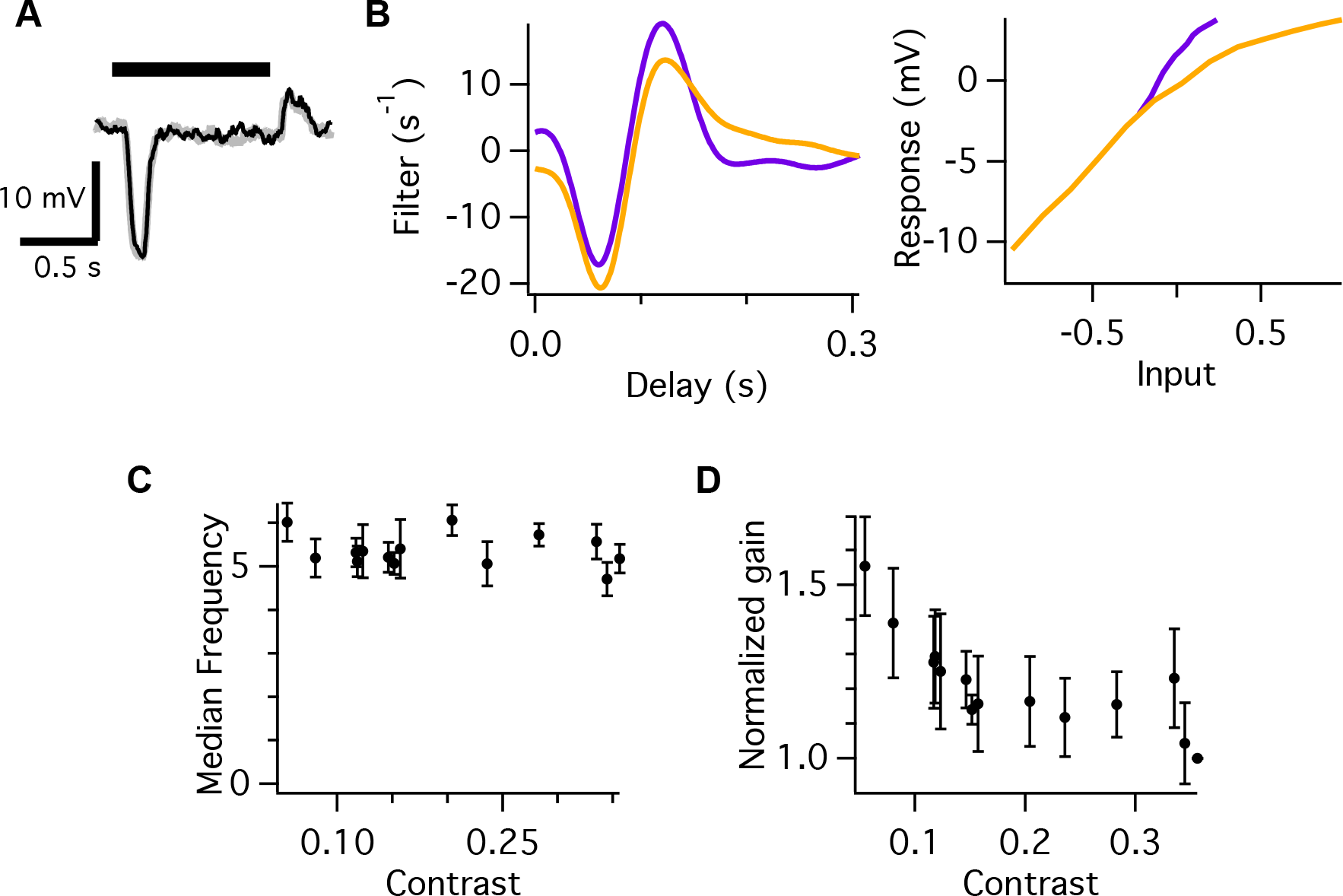
Bipolar cells do not change their temporal frequency during contrast adaptation. **A.** Two superimposed traces of the flash response from an example Off bipolar cell. Bar indicates the time of light On. **B.** LN model for bipolar cells responding to a uniform field stimulus as in Fig. 1 that changed every 20 s to a new contrast ranging between 5 – 35 %. **C.** Median temporal frequency computed from the linear temporal filter, as a function of contrast, averaged over all cells. **D.** Normalized gain as a function of contrast, averaged over six bipolar cells.

## References

1. Shapley RM, Victor JD (1978) The effect of contrast on the transfer properties of cat retinal ganglion cells. J Physiol (Lond) 285:275–298.

2. Nagel KI, Doupe AJ (2006) Temporal processing and adaptation in the songbird auditory forebrain. Neuron 51(6):845–859.

3. Laughlin S (1981) A simple coding procedure enhances a neuron's information capacity. Z Naturforsch, C, Biosci 36(9-10):910–912.

4. Fairhall AL, Lewen GD, Bialek W, de Ruyter Van Steveninck RR (2001) Efficiency and ambiguity in an adaptive neural code. Nature 412(6849):787–792.

5. Atick J (1992) Could information theory provide an ecological theory of sensory processing? Network: Computation in neural systems.

6. Baccus SA (2007) Timing and computation in inner retinal circuitry. Annu Rev Physiol 69:271–290.

7. Baccus SA, Meister M (2002) Fast and slow contrast adaptation in retinal circuitry. Neuron 36(5):909–919.

8. Singh NC, Theunissen FE (2003) Modulation spectra of natural sounds and ethological theories of auditory processing. J Acoust Soc Am 114(6 Pt 1):3394–3411.

9. Van Hateren J (1993) Spatiotemporal contrast sensitivity of early vision. Vision Res.

10. Barlow HB, Fitzhugh R, Kuffler SW (1957) Change of organization in the receptive fields of the cat's retina during dark adaptation. J Physiol (Lond) 137(3):338–354.

11. De Valois RL, Morgan H, Snodderly DM (1974) Psychophysical studies of monkey vision. 3. Spatial luminance contrast sensitivity tests of macaque and human observers. Vision Res 14(1):75–81.

12. Enroth-Cugell C, Robson JG (1966) The contrast sensitivity of retinal ganglion cells of the cat. J Physiol (Lond) 187(3):517–552.

13. Beaudoin DL, Borghuis BG, Demb JB (2007) Cellular basis for contrast gain control over the receptive field center of mammalian retinal ganglion cells. J Neurosci 27(10):2636–2645.

14. Stromeyer CF, Martini P (2003) Human temporal impulse response speeds up with increased stimulus contrast. Vision Res 43(3):285–298.

15. Manookin MB, Demb JB (2006) Presynaptic mechanism for slow contrast adaptation in mammalian retinal ganglion cells. Neuron 50(3):453–464.

16. Jarsky T, et al. (2011) A synaptic mechanism for retinal adaptation to luminance and contrast. J Neurosci 31(30):11003–11015.

17. Chance FS, Nelson SB, Abbott LF (1998) Synaptic depression and the temporal response characteristics of V1 cells. J Neurosci 18(12):4785–4799.

18. Ozuysal Y, Baccus SA (2012) Linking the computational structure of variance adaptation to biophysical mechanisms. Neuron 73(5):1002–1015.

19. Nagel KI, Wilson RI (2011) Biophysical mechanisms underlying olfactory receptor neuron dynamics. Nat Neurosci 14(2):208–216.

20. Friedlander T, Brenner N (2009) Adaptive response by state-dependent inactivation. Proc Natl Acad Sci USA 106(52):22558–22563.

21. Gaudry KS, Reinagel P (2007) Contrast adaptation in a nonadapting LGN model. J Neurophysiol 98(3):1287–1296.

22. Cui Y, Wang YV, Park SJH, Demb JB, Butts DA (2016) Divisive suppression explains high-precision firing and contrast adaptation in retinal ganglion cells. Elife 5:220.

23. Victor JD (1987) The dynamics of the cat retinal X cell centre. J Physiol (Lond) 386:219–246.

24. Kim KJ, Rieke F (2001) Temporal contrast adaptation in the input and output signals of salamander retinal ganglion cells. J Neurosci 21(1):287–299.

25. Chichilnisky EJ (2001) A simple white noise analysis of neuronal light responses. Network 12(2):199–213.

26. Rieke F (2001) Temporal contrast adaptation in salamander bipolar cells. J Neurosci 21(23):9445–9454.

27. Werblin FS (2010) Six different roles for crossover inhibition in the retina: correcting the nonlinearities of synaptic transmission. Vis Neurosci 27(1–2): 1–8.

28. Colquhoun D, Hawkes AG (1977) Relaxation and fluctuations of membrane currents that flow through drug-operated channels. Proc R Soc Lond, B, Biol Sci 199(1135):231–262.

29. Hodgkin AL, Huxley AF (1952) A quantitative description of membrane current and its application to conduction and excitation in nerve. J Physiol (Lond) 117(4):500–544.

30. Burrone J, Lagnado L (2000) Synaptic depression and the kinetics of exocytosis in retinal bipolar cells. J Neurosci 20(2):568–578.

31. Gomis A, Burrone J, Lagnado L (1999) Two actions of calcium regulate the supply of releasable vesicles at the ribbon synapse of retinal bipolar cells. J Neurosci 19(15):6309–6317.

32. Neves G, Lagnado L (1999) The kinetics of exocytosis and endocytosis in the synaptic terminal of goldfish retinal bipolar cells. J Physiol (Lond) 515 (Pt 1):181–202.

33. Geffen MN, de Vries SEJ, Meister M (2007) Retinal ganglion cells can rapidly change polarity from Off to On. PLoS Biol 5(3):e65.

34. Zaghloul KA, Boahen K, Demb JB (2003) Different circuits for ON and OFF retinal ganglion cells cause different contrast sensitivities. J Neurosci 23(7):2645–2654.

35. Chichilnisky EJ, Kalmar RS (2002) Functional asymmetries in ON and OFF ganglion cells of primate retina. J Neurosci 22(7):2737–2747.

36. Kastner DB, Baccus SA (2013) Spatial segregation of adaptation and predictive sensitization in retinal ganglion cells. Neuron 79(3):541–554.

37. Kastner DB, Baccus SA, Sharpee TO (2015) Critical and maximally informative encoding between neural populations in the retina. Proc Natl Acad Sci USA 112(8):2533–2538.

38. Molnar A, Hsueh H-A, Roska B, Werblin FS (2009) Crossover inhibition in the retina: circuitry that compensates for nonlinear rectifying synaptic transmission. J Comput Neurosci 27(3):569–590.

39. Liang Z, Freed MA (2010) The ON pathway rectifies the OFF pathway of the mammalian retina. J Neurosci 30(16):5533–5543.

40. Pang J-J, Gao F, Wu SM (2007) Cross-talk between ON and OFF channels in the salamander retina: indirect bipolar cell inputs to ON-OFF ganglion cells. Vision Res 47(3):384–392.

41. Zaghloul KA, Boahen K, Demb JB (2005) Contrast adaptation in subthreshold and spiking responses of mammalian Y-type retinal ganglion cells. J Neurosci 25(4):860–868.

42. Chander D, Chichilnisky EJ (2001) Adaptation to temporal contrast in primate and salamander retina. J Neurosci 21(24):9904–9916.

43. Liu JK, Gollisch T (2015) Spike-Triggered Covariance Analysis Reveals Phenomenological Diversity of Contrast Adaptation in the Retina. PLoS Comput Biol 11(7):e1004425.

44. Behnia R, Clark DA, Carter AG, Clandinin TR, Desplan C (2014) Processing properties of ON and OFF pathways for Drosophila motion detection. Nature 512(7515):427–430.

45. Pang J-J, Gao F, Wu SM (2004) Stratum-by-stratum projection of light response attributes by retinal bipolar cells of Ambystoma. J Physiol (Lond) 558(Pt 1):249–262.

46. Franke K, et al. (2017) Inhibition decorrelates visual feature representations in the inner retina. Nature 542(7642):439–444.

47. Baylor DA, Hodgkin AL (1974) Changes in time scale and sensitivity in turtle photoreceptors. J Physiol (Lond) 242(3):729–758.

48. Pugh EN, Nikonov S, Lamb TD (1999) Molecular mechanisms of vertebrate photoreceptor light adaptation. Current Opinion in Neurobiology 9(4):410–418.

49. Weckström M, Laughlin SB (1995) Visual ecology and voltage-gated ion channels in insect photoreceptors. Trends Neurosci 18(1): 17–21.

50. Pan ZH (2000) Differential expression of high-and two types of low-voltage-activated calcium currents in rod and cone bipolar cells of the rat retina. J Neurophysiol 83(1):513–527.

51. Pan ZH (2001) Voltage-activated Ca2+ channels and ionotropic GABA receptors localized at axon terminals of mammalian retinal bipolar cells. Vis Neurosci 18(2):279–288.

52. Sharpee TO, Nagel KI, Doupe AJ (2011) Two-dimensional adaptation in the auditory forebrain. J Neurophysiol 106(4): 1841–1861.

53. Kastner DB, Baccus SA (2011) Coordinated dynamic encoding in the retina using opposing forms of plasticity. Nat Neurosci 14(10):1317–1322.

54. Kastner DB, Baccus SA (2011) Coordinated dynamic encoding in the retina using opposing forms of plasticity. Nat Neurosci 14(10):1317–1322.

55. Smirnakis SM, Berry MJ, Warland DK, Bialek W, Meister M (1997) Adaptation of retinal processing to image contrast and spatial scale. Nature 386(6620):69–73.

56. Korenberg MJ, Hunter IW (1986) The identification of nonlinear biological systems: LNL cascade models. Biol Cybern 55(2-3): 125–134.

